# Solving the trade-off by differences in handling of intracellular K^+^: why substrate translocation by the dopamine transporter but not by the serotonin transporter is voltage-dependent

**DOI:** 10.1101/2020.07.09.196642

**Authors:** Shreyas Bhat, Marco Niello, Klaus Schicker, Christian Pifl, Harald H. Sitte, Michael Freissmuth, Walter Sandtner

## Abstract

The dopamine transporter (DAT) retrieves dopamine into presynaptic terminals after synaptic release. The concentrative power of DAT is thought to be fueled by the transmembrane Na^+^ gradient, but it is conceivable that DAT can also rely on other energy sources, e.g. membrane voltage and/or the K^+^ gradient. Here, we recorded uptake of dopamine or the fluorescent substrate APP^+^ ((4-(4-dimethylamino)phenyl-1-methylpyridinium) in DAT-expressing cells under voltage control. We show that DAT differs substantially from the closely related serotonin transporter (SERT): substrate uptake by DAT was voltage-dependent, intracellular K^+^ binding to DAT was electrogenic but transient in nature thus precluding antiport of K^+^ by DAT. There is a trade-off between maintaining constant uptake and harvesting membrane potential for concentrative power. Based on our observations, we conclude that subtle differences in the kinetics of co-substrate ion binding allow closely related transporters to select between voltage-independent uptake and high concentrative power.

## Introduction

The serotonin transporter (SERT) and the dopamine transporter (DAT) are members of the solute carrier 6 family (SLC6). In the brain, SERT and DAT mediate reuptake of released serotonin and dopamine, respectively, into the presynaptic specialization neurons (*Kristensen et al., 2011*). By this action, they terminate monoaminergic signaling and - in concert with the vesicular monoamine transporters - replenish vesicular stores. The two transporters are secondary active in nature; they utilize the free energy contained in the transmembrane Na^+^ gradient (established by the Na^+^/K^+^ pump) to drive concentrative monoamine uptake into cells in which they are expressed (*Burtscher et al., 2019*). SLC6 transporters are believed to operate via the alternate access mechanism wherein they endure a closed loop of partial reactions constituting a complete transport cycle (*Jardetzky, 1966; Rudnick and Sandtner, 2019*). These partial reactions require conformational rearrangements and binding/unbinding reactions of substrate and co-substrate ions. It is gratifying to note that crystal structures obtained from the prokaryotic homolog LeuT, drosophila DAT and human SERT itself support the general concept of alternate access (*Yamashita et al., 2005; Penmatsa et al., 2013; Coleman et al., 2016*). These crystal structures also reveal SERT and DAT to be closely related structurally. This is evident from the root mean square deviation, which differs by only approximately 1 Å between the outward facing structure of human SERT and drosophila DAT. SERT and DAT also share a rich and partially overlapping pharmacology (*Sitte and Freissmuth, 2015)*: there are many inhibitors, which are used for the treatment of neuropsychiatric disorders (major depression, general anxiety disorder and attention-deficit hyperactivity disorder), are highly selective for either transporter, but many illicit drugs, which induce reverse transport, target both, DAT and SERT (*Hasenhuetl et al., 2019; Niello et al., 2020*).

Despite their similarity in structure and function, the two transporters differ in many more aspects than just ligand recognition: the transport stoichiometry of SERT and DAT is considered to be electroneutral and electrogenic, respectively. It has long been known that SERT antiports intracellular K^+^ (*Rudnick and Nelson, 1978*); for DAT, on the other hand, intracellular K^+^ is thought to be immaterial (*Sonders et al., 1997; Erreger et al., 2008*). If true, only SERT can utilize the chemical potential of the cellular K^+^ gradient to establish and maintain a substrate gradient. It has therefore remained enigmatic, why closely related transporters can differ so fundamentally in their stoichiometry and their kinetic decision points. In this context, it is worth pointing out that the effects of intracellular K^+^ (K_in_^+^), intracellular Na^+^ (Na_in_^+^) and membrane voltage on the transport cycle of SERT have been recently analyzed in great detail (*Hasenhuetl et al., 2016*). However, much less is known on how these factors definitively impinge on the transport cycle of DAT (*Sonders et al., 1997; Hoffman et al., 1999; Prasad and Amara, 2001; Erreger et al., 2008; Li et al., 2015*). In this study, we investigated the role of intracellular K^+^ and voltage on substrate transport through DAT. To this end, we coupled direct measurements of substrate induced currents with simultaneous recordings of uptake of the fluorescent substrate APP^+^ (4-(4-dimethylamino)phenyl-1-methylpyridinium) into single HEK293 cells expressing DAT under voltage control. These measurements were conducted in the whole cell patch clamp configuration, which allowed for control of the intra- and extracellular ion composition via the electrode and bath solution, respectively. Our analysis revealed that K_in_^+^ did bind to the inward facing state of DAT but - in contrast to SERT - K_in_^+^ was released prior the return step from the substrate free inward to the substrate free outward facing conformations. We also found that substrate uptake by DAT, unlike SERT, was voltage-dependent in the presence of K_in_^+^. Moreover, the absence of K_in_^+^ had no appreciable effect on the transport rate of DAT. The transient nature of K_in_^+^ binding was incorporated into a refined kinetic model, which highlights the differences between SERT and DAT. Notably, this model allows for a unifying description, which attributes all existing functional differences between SERT and DAT to the difference in the handling of K_in_^+^.

### Experimental procedures

#### Cell culture

Human embryonic kidney 293 (HEK293) cells were grown in Dulbecco’s Modified Eagle Medium, 10% heat-inactivated fetal bovine serum and 50 mg/l gentamicin on 60 or 100 mm tissue culture dishes (Falcon) at 37°C and 5% CO_2_/95% air. For stable expression of human DAT in HEK cells the expression vector pRc/CMV was used as described previously (*Sitte et al., 1998*).

#### Whole cell patch clamping

Whole cell patch clamp experiments were performed on HEK293 cells stably expressing DAT. Twenty-four hours prior to patching, the cells were seeded at low density on PDL coated 35 mm plates. Substrate-induced DAT currents were recorded under voltage clamp. Cells were continuously superfused with a physiological external solution that contains 140 mM NaCl, 2.5 mM CaCl_2_, 2 mM MgCl_2_, 20 mM glucose, and 10 mM HEPES (pH adjusted to 7.4 with NaOH, final Na^+^ concentration was 146 mM). Pipette solution mimicking the internal ionic composition of a cell (referred to as normal internal solution henceforth, used in Fig.1, Fig. 2, Fig. 5D) contained 133 mM potassium gluconate, 6 mM NaCl, 1 mM CaCl_2_, 0.7 mM MgCl_2_, 10 mM HEPES, 10 mM EGTA (pH adjusted to 7.2 with KOH, final K_in_^+^ concentration 160 mM). A low Cl^-^ internal solution (Fig. 5G, 5H-final Cl^-^ concentration 0.5 mM) was made by replacing CaCl_2_ and MgCl_2_ in normal internal solution by CaMES_2_ (MES - 2-(N-morpholino)ethanesulfonic acid) and Mg-Acetate, respectively and 5.5 mM of NaCl with NaMES. A high Cl^-^ internal solution was made by replacing potassium gluconate in the normal internal solution with KCl (Fig. 5G, 5H – final Cl^-^ concentration 142.4 mM). A Na_in_^+^ and K_in_^+^-free internal solution (Fig. 3D, Fig. 3F, Fig. 4A-G, Fig. 5E) was made by replacing NaCl and potassium gluconate in the normal internal solution with equimolar concentrations of NMDG chloride (titrated to pH of 7.2 using NMDG). A Na_in_^+^-free 160 mM K_in_^+^ internal solution (Fig.4B, Fig.4D, Fig. 4E, Fig.4F, Fig.5A-B, Fig.5F) was made by replacing NaCl with NaMES (titrated to pH of 7.2 with KOH; final K^+^ concentration 160 mM). An internal solution with high NaCl (used in Fig. 3A-C, Fig.3E, Fig.3F) was made by replacing potassium gluconate of the normal internal solution with equimolar concentration of NaCl (pH adjusted to 7.2 with NaOH; final Na^+^ concentration 160 mM). A high Li^+^ internal solution was made by replacing potassium gluconate in the normal internal solution with 133 mM of LiCl (pH adjusted to 7.2 with LiOH, Fig. 5C; final Li^+^ concentration 160 mM). Currents elicited by dopamine or APP^+^, a fluorescent substrate of DAT (IDT307, Sigma Aldrich), were measured at room temperature (20-24^0^C) using an Axopatch 200B amplifier and pClamp 10.2 software (MDS Analytical Technologies). Dopamine or APP^+^ was applied using a DAD-12 superfusion system and a 4-tube perfusion manifold (ALA Scientific Instruments), which allowed for rapid solution exchange. Current traces were filtered at 1 kHz and digitized at 10 kHz using a Digidata 1550 (MDS Analytical Technologies). Current amplitudes and accompanying kinetics in response to substrate application were quantified using Clampfit 10.2 software (Molecular Devices). Passive holding currents were subtracted, and the traces were filtered using a 100-Hz digital Gaussian low-pass filter.

**Fig. 1.**
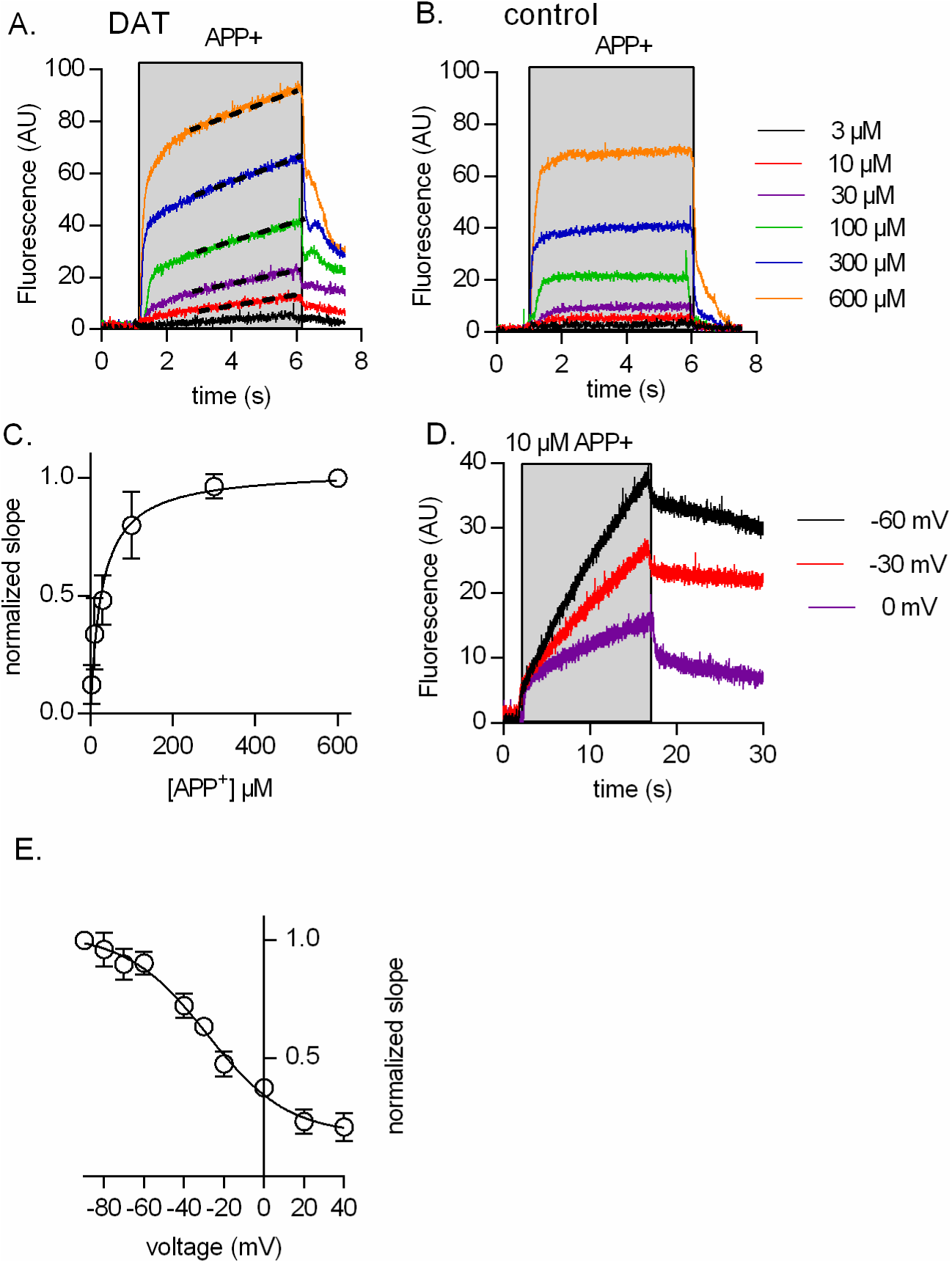
Single cell uptake of APP^+^. **A & B.** Representative traces of APP^+^ fluorescence emission recorded from a cell expressing DAT (A) and a control cell (B) Cells were held at - 60 mV in the whole-cell patch-clamp configuration. The extra- and intracellular solution contained physiological ion concentrations. APP^+^ was applied for a period 5 seconds at the indicated concentrations. The dashed black lines in panel A are the linear regression to the continuous rise in APP^+^ fluorescence, which is absent in panel B. The fast initial rise seen in A. and B. reflects reversible APP^+^ binding to the plasma membrane. **C.** The slopes obtained from the linear regression (as shown in panel A) were normalized to the slope recorded at 600 µM APP^+^ to account for inter-cell variation and plotted as function of APP^+^ concentration (n = 8). The solid black line indicates a fit of the Michaelis Menten equation to the data points (open circles; error bars are SD). The K_M_ was 27.7 µM (95% confidence interval: 20.6 - 34.9 µM). **D.** Representative traces showing APP^+^ uptake recorded from an DAT expressing cell at - 60 mV, - 30 mV and 0 mV, respectively. The cell was exposed to 10 µM APP^+^ for a period of 15 seconds. **E.** Normalized slopes at 10 µM APP^+^ as a function of voltage (range: - 90 mV to + 40 mV). The rate of fluorescence increase was normalized (- 90 mV = 1) to account for inter-cell variation. The solid black line was drawn by fitting the data points to the Boltzmann equation (n = 12). The estimated valence and V_50_ were 1.3 (95% confidence interval: 0.9 - 2.2) and - 28.5mV (95% confidence interval: - 35 to - 21.9 mV], respectively.

**Fig. 2.**
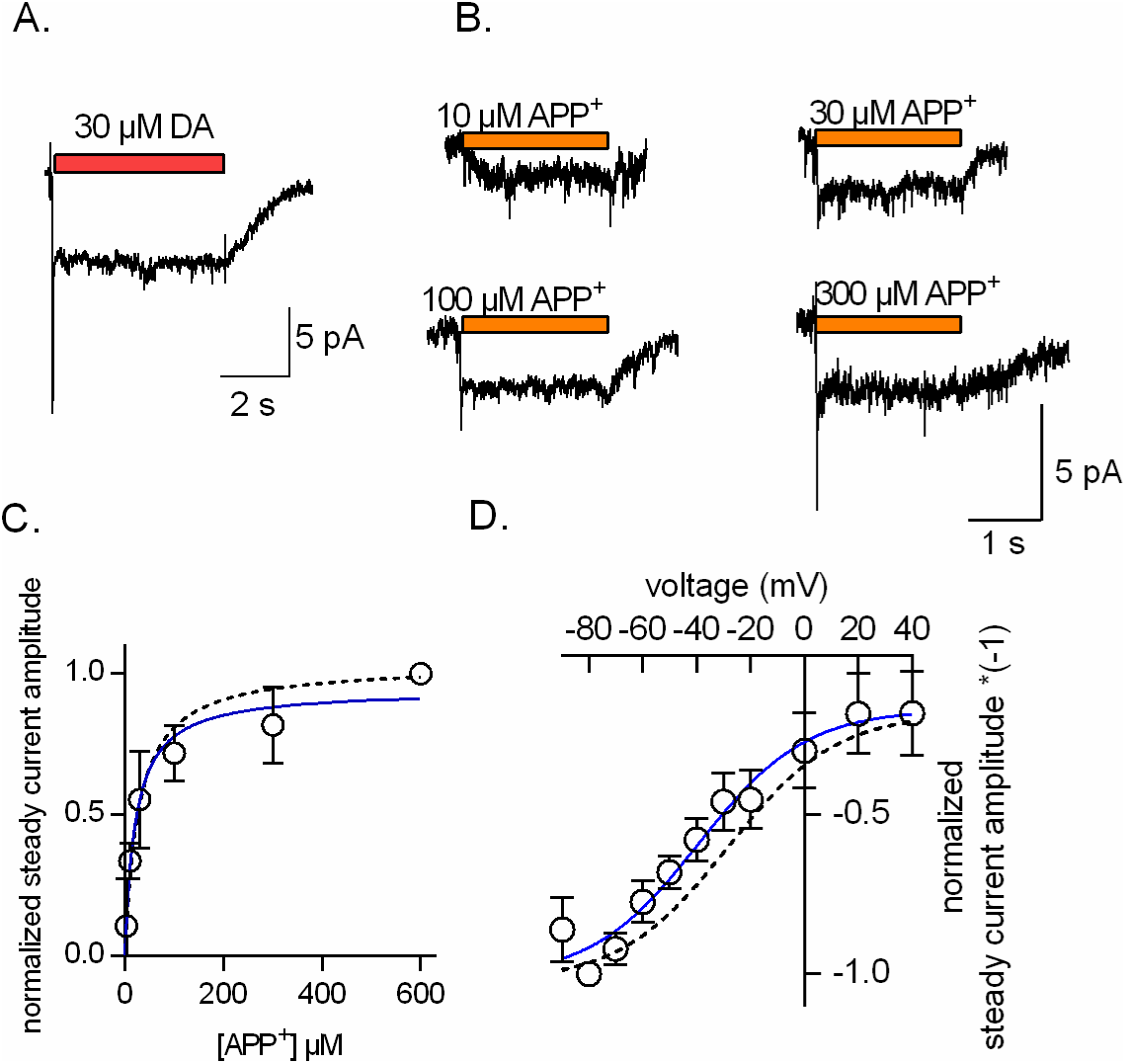
APP^+^ induced currents through DAT. **A.** Representative trace of a dopamine-induced current through DAT. The cell was clamped to - 60 mV and 30 µM dopamine was applied for a period of 5 seconds. The current features a peak and a steady component. **B.** Representative traces of currents elicited by 10 µM, 30 µM, 100 µM and 300 µM APP^+^. At concentrations higher than 10 µM the currents displayed a peak and a steady component. **C.** Normalized steady current amplitude as a function of APP^+^ concentration (open circles, n = 6; error bars are SD). The current amplitude recorded at 600 µM APP^+^ was set 1.0 to normalize for inter-cell variation. The solid blue line was drawn by fitting the data points to the Michaelis-Menten equation. The K_M_ was 21.5 µM (95% confidence interval: 13 - 30 µM). The dashed black line is the fit through the data in Fig. 1C (i.e. the slopes of APP^+^ fluorescence). **D.** Voltage dependence of the normalized steady current amplitude (multiplied by - 1) evoked by 600 µM APP^+^ (open circles; n = 6; error bars are SD). The solid black line is the fit of the Boltzmann equation to the data points (valence: 1.3; 95% confidence interval: 0.72 - 2.4; V_50_: - 39.3 mV; 95% confidence interval: - 53.9 **-** to 24.7 mV). The dashed line is the sign-inverted fit to the data shown in Fig. 1E (i.e. the slopes of APP^+^ fluorescence).

**Fig. 3.**
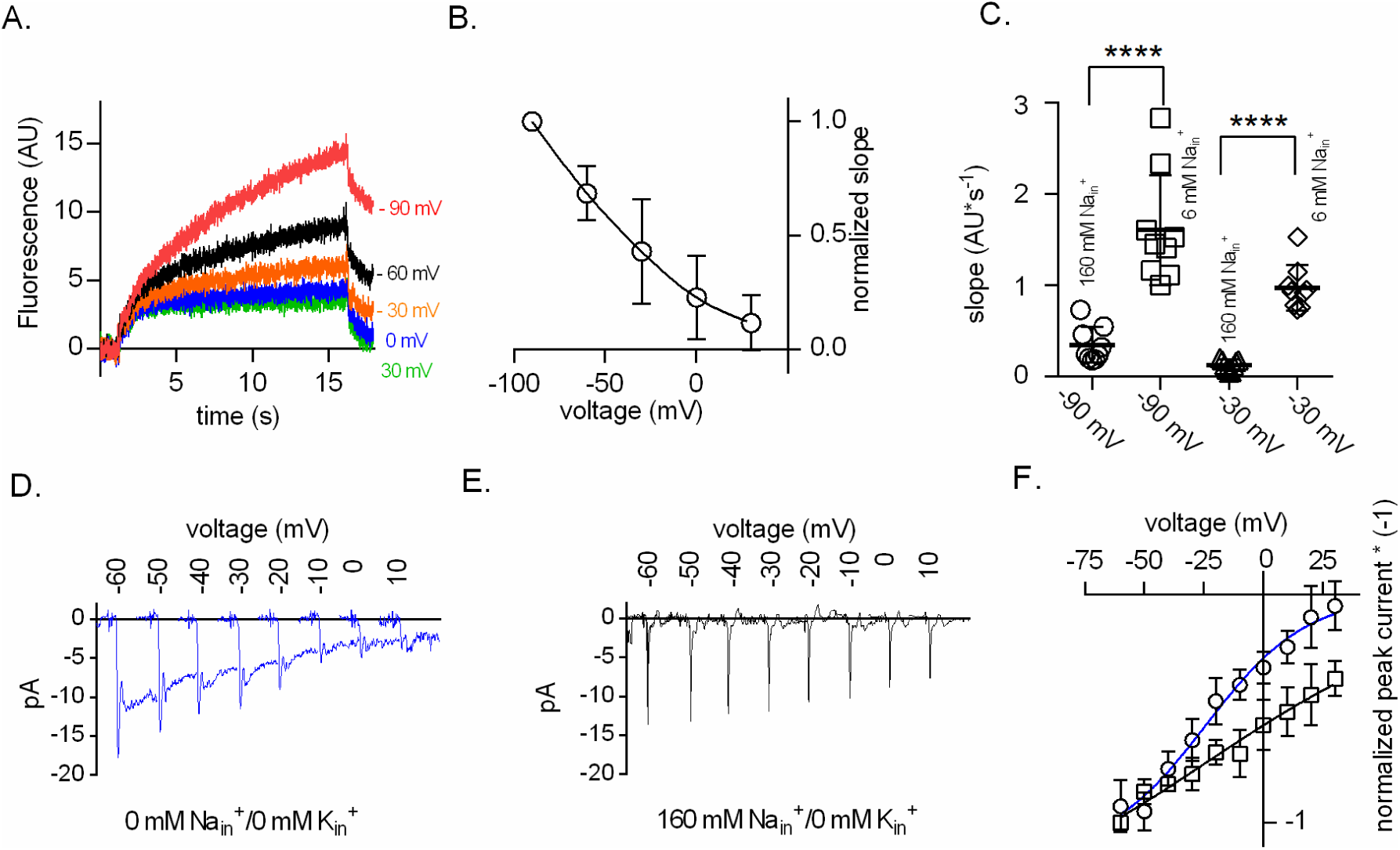
Na^+^ binding to the inward-facing conformation of DAT is voltage-dependent. **A.** Representative recordings of APP^+^ uptake into a cell expressing DAT. The intracellular solution in this experiment contained 160 mM Na^+^. The cell was repeatedly exposed to 10 µM APP^+^ for 15 seconds. The displayed traces were recorded at - 90 mV, - 60 mV, - 30 mV, 0 mV and + 30 mV, respectively. **B.** Normalized slopes of the APP^+^ induced rise in fluorescence plotted as a function of voltage (open circles; n = 9; error bars are SD). The solid black line is the line fit of the data points to the Boltzmann equation. **C.** Comparison of the slopes of APP^+^ fluorescence in 9 cells each measured at 6 mM Na_in_^+^ and at 160 Na_in_^+^. The two groups of cells were compared at - 90 mV and - 30 mV. Open circles (160 mM Na_in_^+^, - 90 mV), open squares (6 mM Na_in_^+^, - 90 mV), open triangles (160 mM Na_in_^+^, - 30 mV) and open diamonds (6 mM Na_in_^+^, - 30 mV). The high intracellular Na^+^ concentration was inhibitory at both potentials (p < 0.0001); Mann Whitney test. **D.** Representative traces of currents elicited by dopamine (30 µM) recorded at potentials between - 60 mV and + 10 mV. Shown are the first 200 ms of the recording at the indicated potentials. In the experiment Na_in_^+^ and K_in_^+^ were replaced by the inert cation NMDG^+^. Under these conditions, the current featured a peak and a steady component. **E.** Representative traces obtained as in panel D but with 160 mM Na_in_^+^. Under these conditions, the peak current was present but the steady current component was suppressed. **F.** Normalized peak current amplitudes (multiplied by - 1) as a function of voltage. Plotted in the same graph are the data for zero Na_in_^+^/ K_in_^+^ (open circles; n = 6) and for 160 mM Na_in_^+^ (open squares; n = 6), respectively. The solid blue and black lines were drawn by fitting the data to the Boltzmann equation. In the absence of intracellular Na^+^ and K^+^ the estimated valence was 1.22 (95% confidence interval: 0.73 - 3.65) and V_50_ = - 24.5 mV (95% confidence interval: −36.5 - −12.5 mV). In the presence of 160 mM Na_in_^+^, the valence was 0.39 (95% confidence interval: 0.13 – 2). We obtained the latter estimate by imposing the following constraints onto the fit: (i) top value < zero, (ii) shared V_50_ between data sets.

**Fig. 4.**
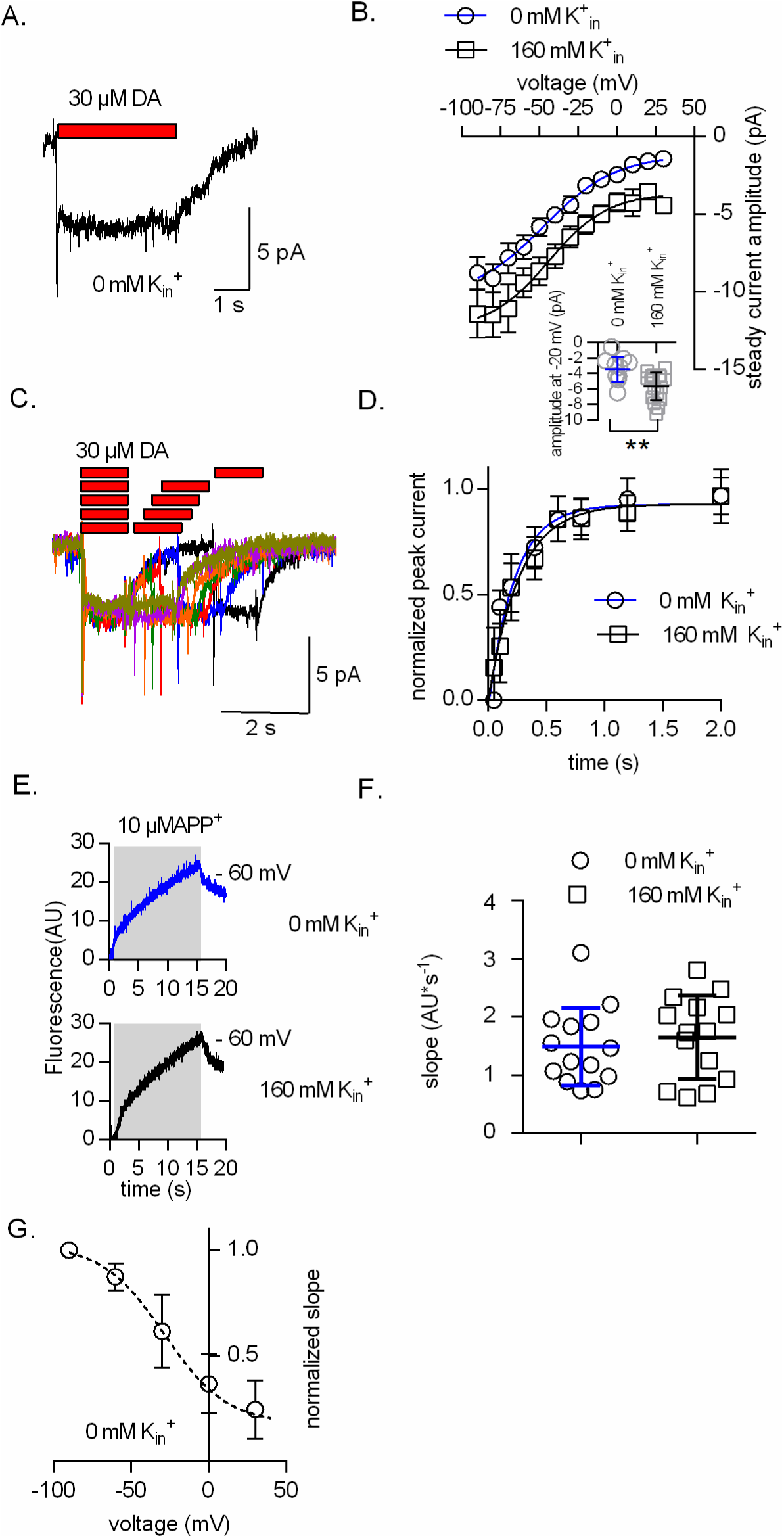
Effects of intracellular K^+^ on dopamine-induced currents and APP^+^ uptake through DAT. **A.** representative trace of a current recorded from a cell expressing DAT, which was held at - 60 mV. The current was evoked by a 5 second application of 30 µM dopamine. The intracellular solution was devoid of K^+^. **B.** Comparison of the steady current amplitudes in the absence (open circles) and presence of K_in_^+^ (open squares) (n = 9 each). The currents were recorded in a voltage range from - 90 mV to + 30 mV. The solid blue and black line were drawn by fitting the data points to the Boltzmann equation. K_in_^+^ (0 mM): valence = 1.03 (95% confidence interval: 0.46 - 4.8]; K_in_^+^ (160 mM): valence = 1.02 (95% confidence interval: 0.40 - 1.9). The inset shows the comparison of amplitudes recorded at - 20 mV. The difference in amplitude at this potential was significant (p = 0.003, Mann-Whitney test). The error bars in the graph are SEM, but SD in the inset. **C.** The traces are an example of a two-pulse protocol, which is schematically outlined above the traces: all currents were evoked by the application of 30 µM dopamine. The reference pulse (first red bar giving rise to the first current) was followed by a test pulse (second red bar giving rise to the second current). The time intervals (indicated by the gap) between the two pulses were 0.1 s, 0.2 s, 0.4 s, 0.6 s, 0.8 s, 1.2 s and 2 s. **D.** Normalized peak currents as a function of the time interval between reference and test pulse for 0 mM K_in_^+^ (open circles; n = 6; error bars are SD) and 160 mM K_in_^+^ (open squares; n = 6; error bars are SD). The currents were recorded at - 50 mV. The solid blue and black lines were drawn by fitting the data points to mono-exponential rises. Tau (0 mM K_in_^+^) = 0.22 s (95% confidence interval: 0.16-0.40 s); Tau (160 mM K_in_^+^) = 0.26 s (95% confidence interval: 0.19 - 0.46 s). The two time courses were not different (p=0.86; F-test). **E.** Representative traces of recordings of APP^+^ uptake (0 mM K^+^ upper panel; 160 mM K^+^ lower panel) from two different cells expressing DAT. The cells were exposed to 10 µM APP^+^ for 15 seconds at a holding potential of - 60 mV. F. Comparison of the slopes for the two conditions (open circles – 0 mM K_in_^+^; open squares-160 mM K_in_^+^) (n = 14; each). The difference in APP^+^ uptake between the two groups were not different (p = 0.53; Mann-Whitney test). G. Slope of APP^+^ fluorescence measured at - 90 mV, - 60 mV, - 30 mV, 0 mV and + 30 mV in the absence of K_in_^+^ (open circles- n = 8; error bars are SD). The slopes were normalized to the slope at - 90 mV. For comparison we show the fit to the data in Fig.1E (160 mM K_in_^+^ – black dashed line).

**Fig. 5.**
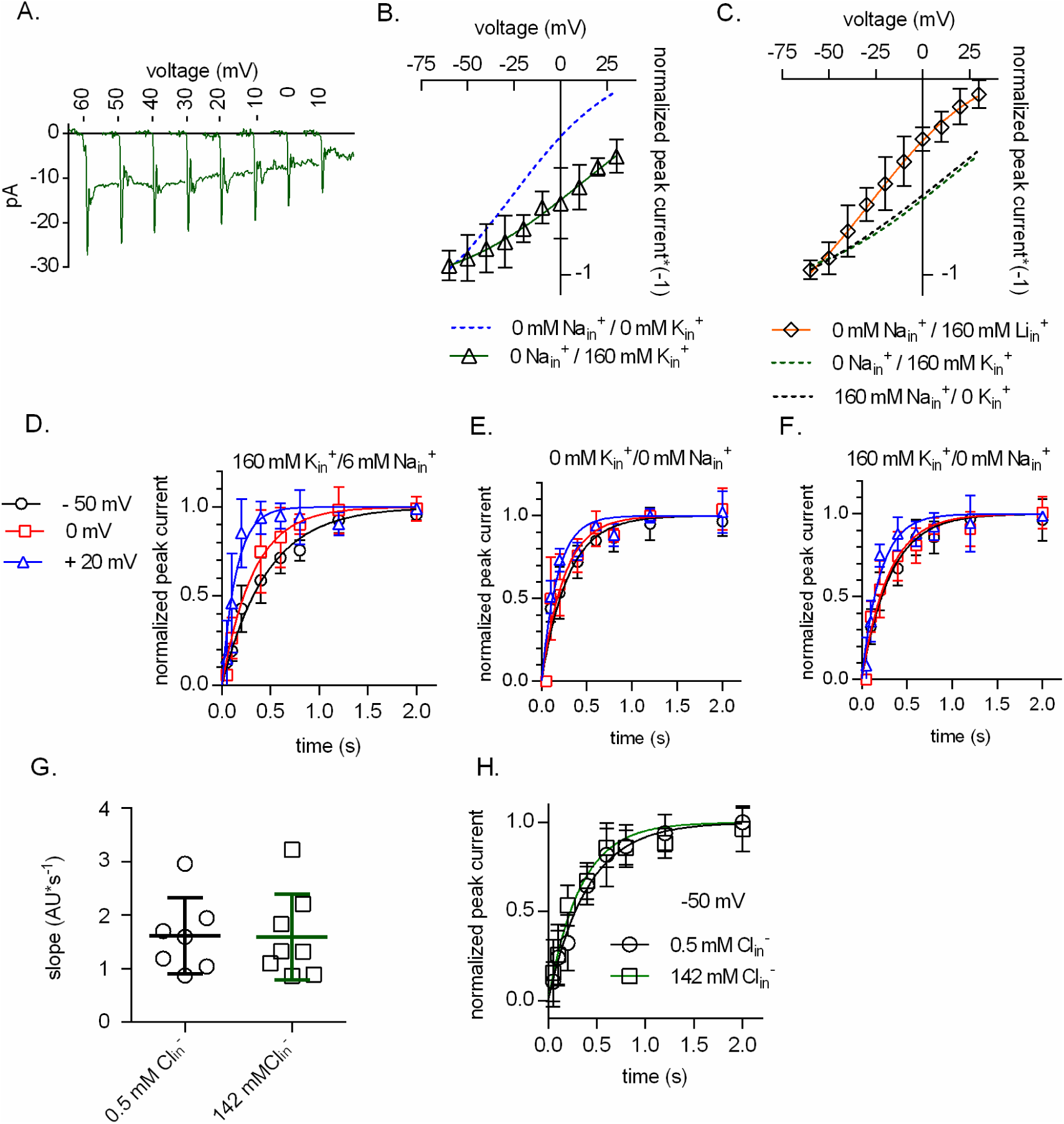
Binding of K^+^ to the inward facing conformation of DAT is voltage-dependent. **A.** Representative traces of dopamine-induced currents through DAT measured in the voltage range from - 60 mV to + 10 mV. The currents were evoked with 30 µM dopamine. The intracellular solution contained 160 mM K^+^. The traces show the first 200 ms of the current at the indicated potentials. **B.** Normalized peak currents in the presence of 160 mM K_in_^+^ (open triangles; n = 6) as a function of voltage. The green line is a Boltzmann fit through the data points. The valence was 0.38 (95% confidence interval: 0.05 – 2). We obtained this estimate imposing the following constraints onto the fit: (i) top value < zero; (ii) shared V_50_ between data sets. The dashed blue line (i.e. fit from the data in Fig. 3F - 0 mM Na_in_^+^/ K_in_^+^) was added for comparison. **C.** Normalized peak current amplitude as a function of voltage with 160 mM Li^+^ present intracellularly (open diamonds; n = 6). The red solid line is the fit of the data to the Boltzmann equation. The valence was 1.23 (95% confidence interval: 0.60 - 2.41) and V_50_ = - 24.5 mV (95% confidence interval: - 36.5 - - 12.5 mV). The dashed black and green lines are the fits for 152 mM Na_in_^+^ and 160 mM K_in_^+^, respectively (*cf*. Fig. 3F and Fig. 5B). **D. E.** and **F.** are the results obtained from applying “two pulse protocols” at - 50 mV (black open circles), 0 mV (red open squares) and + 20 mV (blue open triangles). The tested conditions were 160 mM K_in_^+^/6 mM Na_in_^+^ in D., 0 mM K_in_^+^/0 mM Na_in_^+^ in E. and 160 mM K_in_^+^/0 mM Na_in_^+^ in F. (n = 6; each). The solid lines in black, red and blue were drawn by fitting the data to the equation for a mono-exponential rise. The rates in D. were 2.16 s^-1^ (95% confidence interval: 1.93 - 2.39 s^-1^), 3.08 s^-1^ (95% confidence interval: 2.39-3.78 s^-1^) and 6.77 s^-1^ (95% confidence interval: 4.60 - 8.93 s^-1^) for - 50 mV, 0 mV and + 20 mV, respectively. The rates in E. were 3.40 s^-1^ (95% confidence interval: 2.79 - 4.01 s^-1^), 3.99 s^-1^ (95% confidence interval: 3.05 - 4.93 s^-1^) and 5.71 s^-1^ (95% confidence interval: 4.47 - 6.95 s^-1^) for - 50 mV, 0 mV and + 20 mV, respectively. The rates in F. were 3.14 s^-1^ (95% confidence interval: 2.62 - 3.66 s^-1^), 3.38 s^-1^ (95% confidence interval: 2.73- 4.03 s^-1^) and 4.92 s^-1^ (95% confidence interval: 3.80 - 6.04 s^-1^) for - 50 mV, 0 mV and + 20 mV, respectively. G. Comparison of APP^+^ uptake (10 µM) at a low and a high Cl_in_^-^ concentration. Shown are the slopes measured with 0.5 mM Cl_in_^-^ (open circles; n = 7) and with 142 mM Cl_in_^-^ (open squares; n = 8) at - 60 mV. The slopes were not different (p = 0.93; Mann Whitney test) H. Peak current recovery at – 50 mV. Shown are normalized peak current amplitudes as a function of the time interval between the reference and the test pulse. Currents were elicited by rapid application of 30 µM dopamine. Open circles (n = 6) show the peak current recovery with 0.5 mM Cl_in_^-^ and open squares (n = 6) the peak current recovery with 142 mM Cl_in_^-^. The solid black and green line indicate fits of a mono-exponential function to the data points. With 0.5 mM Cl_in_^-^ the recovery rate was 2.49 s^-1^ (95% confidence interval: 2.14 – 2.83 s^-1^); with 142 mM Cl_in_^-^ the recovery rate was 3.04 s^-1^ (95% confidence interval: 2.49 – 2.59 s^-1^). Difference in these rates was not statistically significant (p= 0.086; F-test).

### Fluorescence recordings

Twenty-four hours prior to fluorescence recording, HEK293 cells stably expressing DAT were seeded at low density on PDL-coated 35 mm glass bottom dishes, which have a cover glass (Cellview Cell Culture Dish, Greiner Bio-One GmbH; Germany). On the day of the experiment, individual cells were visualized and patched using a 100x oil-immersion objective. APP^+^, a fluorescent substrate of DAT that has an excitation range from 420-450 nm in polar solvents, was applied to single cells using a perfusion manifold in physiological external solution for 5 seconds at room temperature (20-24°C). APP^+^ uptake into the cell was measured using a LED lamp emitting 440 nm light and a dichroic mirror that reflected the light onto the cells. The emitted fluorescence from the sequestered APP^+^ within the cell was measured using a photomultiplier tube (PMT2102, Thorlabs, United States) mounted on the microscope after it had passed an emission filter (510-530 nm). The signal from the PMT was filtered at 3 kHz, digitized at 10 kHz with an Axon Digidata 1550B and pClamp 10.2 software (MDS Analytical Technologies). The signals (i.e. currents and fluorescence) were acquired with separate channels.

### Kinetic modeling and statistics

The kinetic model for the DAT transport cycle is based on previously reported sequential binding models for DAT (*Erreger et al., 2008*) and SERT (*Hasenhuetl et al., 2016*). State occupancies was calculated by numerical integration of the resulting system of differential equations using the Systems Biology Toolbox (*Schmidt and Jirstrand, 2006*) and MAT LAB 2017a software (Mathworks). The voltage dependence of individual partial reactions was modeled assuming a symmetric barrier as k_ij_ = k^0^_ij_e^-zQijFV/2RT^, where F = 96,485 C·mol^-1^, R = 8.314 JK^-1^mol^-1^, V is the membrane voltage in volts, and T = 293 K (*Läuger, 1991*).

Coupled membrane currents upon application of substrate were calculated as I = -F×NC/N_A_×ΣSz_Qij_(p_i_k_ij_ - p_j_k_ji_), where z_Qij_ is the net charge transferred during the transition, NC is the number of transporters (4 × 10^6^/cell), and N_A_ = 6.022e^23^/mol. The substrate-induced uncoupled current was modeled as a current through a Na^+^-permeable channel with I = P_o_γNC(V_M_ -V_rev_), where P_o_ corresponds to the occupancy of the channel state, γ is the single-channel conductance of 2.4 pS, V_M_ is the membrane voltage, and V_rev_ is the reversal potential of Na^+^ (+ 100 mV). The extracellular and intracellular ion concentrations were set to the values used in the respective experiments. To account for the non-instantaneous onset of the substrate in patch-clamp experiments, we modeled the substrate application as an exponential rise with a time constant of 10 ms. Uptake of APP^+^ was modeled as T_i_ClS*k_off_S_in_ -T_i_Cl*k_on_S_in_*S_in_* NC/NA, where T_i_ClS and T_i_Cl are the respective state occupancies, k_on_S_in_ and k_off_S_in_ are the association and dissociation rate constants of APP^+^ and S_in_ is the intracellular concentration of APP^+^.

Experimental variations are either reported as means ± 95% confidence intervals, means ± SD, or means ± SEM. Current voltage relations were fit to a Boltzmann equation (I = Bottom + ((Bottom-Top) / (1 + exp ((V_50_ -V_M_)/slope))), where I is the current, Bottom and Top the lower level (i.e. maximal current amplitude of the inwardly directed current at negative potentials) and higher level (i.e. maximal current amplitude of the inwardly directed current at positive potentials), respectively and V_50_ the potential at which the current amplitude is half of the span between Bottom and Top. The valence (z) was calculated from the slope: z = (R*T/slope)*F

## Results

### Cellular APP^+^ uptake via DAT is voltage dependent

APP^+^, a fluorescent analog of MPP^+^(1-methy-4-phenylpyridinium), is a substrate of SERT and DAT (*Solis Jr. et al., 2012*). APP^+^ uptake into DAT expressing cells can be monitored with high temporal resolution by the detection of the light emitted by sequestered APP^+^. Here we explored the feasibility of measuring APP^+^ uptake into a single cell under voltage control by employing a photomultiplier tube and a patch clamp amplifier, respectively. In Fig. 1, we show representative traces obtained from a HEK293 cell stably expressing DAT (Fig. 1A) and from an empty HEK293 cell (Fig. 1B). The cells were clamped to - 60 mV in the whole cell configuration and exposed to increasing APP^+^ concentrations (i.e. 3 µM, 10 µM, 30 µM, 100 µM, 300 µM and 600 µM) with a rapid superfusion device for 5 seconds. On APP^+^ application, the fluorescence intensity rose linearly in DAT cells but remained at a constant level in control cells after both cell types show a fast and concentration dependent initial rise in fluorescence. On terminating APP^+^ application, the fluorescence intensity in control cells decayed fully to baseline values. In DAT cells, the fluorescence intensity also initially decreased upon APP^+^ removal, but settled at values above the original baseline. We surmise that the fast rise in fluorescence reports on reversible APP^+^ binding to the plasma membrane. The slow and linear increase in fluorescence, seen only in DAT cells, reflects transporter mediated APP^+^ uptake. Slopes were estimated by linear regression of the slow phase of fluorescence increase in DAT-expressing cells (dashed lines in Fig. 1A). These slopes have the dimension of a rate (i.e. fluorescence*s^-1^) and are hence a suitable readout for the uptake rate of APP^+^. We normalized the slopes to those obtained at the highest concentration of APP^+^ tested (600 µM) to allow for comparison between individual cells. When plotted as a function of the applied APP^+^ concentration, these normalized slopes conformed to the Michaelis-Menten equation (Fig. 1C, solid black line): a K_M_ of 28 µM (95% confidence interval 21 - 35 µM) was estimated by fitting the data to a rectangular hyperbola. We applied APP^+^ at a concentration below K_M_ (10 µM) to DAT-expressing cells for 15 seconds at different membrane voltages.

It is evident from the representative traces shown in Fig. 1D that the slope of fluorescence increase - i.e. the rate of APP^+^ uptake - was highest at - 60 mV and progressively decreased at - 30 mV and 0 mV. In the control cell, changes in voltage did not affect background APP^+^ binding (data not shown). In Fig. 1E, we plotted the normalized slopes induced by 10 µM APP^+^ recorded over a voltage range spanning - 90 mV to + 40 mV. The data show that APP^+^ uptake through DAT was voltage-dependent: the rate of APP^+^ uptake declined at more positive potentials. A fit to the Boltzmann equation (solid line in Fig. 1E) allowed for estimating the valence associated with the transport of APP^+^: we obtained a value of 1.3 (95% confidence interval: 0.9 - 2.2).

### Substrate induced currents via DAT are voltage-dependent

Application of endogenous or synthetic substrates elicits a current through DAT, which can be recorded in the whole-cell patch-clamp configuration (*Sonders et al., 1997; Sitte et al., 1998; Erreger et al., 2008; Li et al., 2015*). Rapid substrate application (i.e. solution exchange rate > 10 s^-1^) allows for resolving two current components in the presence of physiological ionic gradients: (i) an initial peak - and (ii) a steady state current, which are both inwardly directed (*Erreger et al., 2008; Li et al., 2015*). The steady state current persists as long as the substrate is present but deactivates to baseline upon its removal. This is illustrated in Fig. 2A for dopamine, the cognate substrate of DAT, when applied at a saturating concentration (30 µM). APP^+^ also evoked peak and steady-state currents, but the peak current was absent at concentrations below 30 µM (Fig. 2B). This can be rationalized by taking into account that APP^+^ is a low affinity substrate of DAT (see also below, kinetic model section): a K_M_ value of 22 µM (95% confidence interval: 13 - 30 µM) was calculated from the saturation curve (blue line in Fig. 2C). For the sake of comparison, the dashed line in Fig. 2C also shows the saturation curve from Fig. 1C (i.e. normalized slopes of the slow rise in APP^+^ fluorescence): there was an excellent agreement between the APP^+^ uptake and the amplitude of the APP^+^-induced steady current, which confirms that APP^+^-induced currents are transport-associated, that is they reflect the rate of APP^+^ uptake. We note, however, that the average current amplitude at saturation was approximately half of that obtained at a saturating dopamine concentration. We also assessed the voltage-dependence (solid blue line in Fig. 2D) of the steady current amplitude at a saturating APP^+^ (600 µM): the amplitude of the steady currents decreased at positive potentials. The estimated valence for APP^+^ extracted from the fits to the Boltzmann equation was again 1.3. The dashed line in Fig. 2D is the sign-inverted fit to the data shown in Fig. 1E for the rate of APP^+^ uptake. It is evident that two voltage relations are similar.

### Na^+^ binding to the inward facing conformation of DAT is voltage-dependent

Substrate transport through Na^+^ dependent transporters is predicted to be impeded at elevated intracellular Na^+^ concentrations: completion of the transport cycle is contingent on cytosolic release of Na^+^ and substrate, but at high intracellular Na^+^ concentrations the ion binding site will remain occupied due to rebinding of Na^+^. As a consequence, the transporter cannot progress through the transport cycle, which must result in reduced substrate uptake (*Khoshbouei et al., 2003*; *Erreger et al., 2008; Schicker et al., 2012; Hasenhuetl et al., 2015; Hasenhuetl et al., 2016*). We verified this prediction by measuring APP^+^ (10 µM) uptake into DAT expressing cells under voltage control using different intracellular patch pipette solutions. The representative traces in Fig. 3A show uptake of APP^+^ into a cell, in which the concentration of Na_in_^+^ was 160 mM and the voltage was held at - 90 mV, - 60 mV, - 30 mV, 0 mV and 30 mV. Normalized recordings (slope at - 90 mV = 1) from 9 experiments are summarized in Fig. 3B. Surprisingly, we found that APP^+^ was transported by DAT at negative potentials despite the presence of 160 mM Na_in_^+^. In Fig. 3C, we compared the slopes of 9 cells, in which Na_in_^+^ was either 6 mM (i.e., normal internal solution) or 160 mM and the membrane potential was clamped at - 90 mV and - 30 mV. It is evident from this comparison that the high intracellular Na^+^ concentration inhibited APP^+^ uptake at both potentials. However, our data also show that the inhibitory effect of intracellular Na^+^ is lesser at negative voltages (Fig. 3B). This indicates that Na^+^ binding to the inward facing conformation is voltage-dependent, rendering the Na_in_^+^ affinity lower at negative potentials.

We explored this conjecture by measuring the dopamine-induced peak current amplitude of DAT as a function of voltage with intracellular solutions containing either 160 mM Na_in_^+^ or 0 mM Na_in_^+^, respectively. In Fig. 3D and Fig.3E, we show representative current traces for both conditions (i.e. 0 mM Na_in_^+^ in Fig. 3D, 160 mM Na_in_^+^ in Fig.3E). In Fig. 3F, we plotted the normalized peak current voltage relations (n = 6). The blue (0 Na_in_^+^) and black lines (160 mM Na_in_^+^) were drawn by fitting the data points to the Boltzmann equation. The peak current amplitude changed with voltage in both conditions. More importantly, we found the slope of the current voltage relation to be steeper in the absence of Na_in_^+^ (open circles) than in the presence of 160 mM Na_in_^+^ (open squares). This action of Na_in_^+^ can be explained as follows: the transition of substrate and Na^+^-loaded transporter from the outward to the inward facing conformation leads to the opening of the inner vestibule and access to the intracellular bulk solution. This creates an exit path for Na^+^ and hence to its intracellular dissociation. Because the Na^+^ binding site resides within the electric field, the dissociating Na^+^ ions give rise to an inwardly directed current. At 160 mM Na_in_^+^ concentrations, however, Na^+^ is expected to rebind producing an outwardly directed current that reduces the net-charge of the inwardly directed peak current. This results in a diminished voltage dependence. The valences extracted from the Boltzmann fit were 1.22 and 0.39 in 0 mM and 160 mM Na_in_^+^ conditions, respectively. Notably, the difference between these two values was 0.83, which is slightly less than 1. This suggests that DAT releases one Na^+^ ion into the cytosol in each cycle.

### K_in_^+^ mediates uncoupled currents in DAT but does not influence substrate uptake

A recent DAT model considers DAT-mediated currents to be fully coupled to substrate transport (*Erreger et al., 2008*). However, there is experimental evidence suggesting that the dopamine-induced current is, in part, produced by an uncoupled ion flux (*Sonders et al., 1997; Sitte et al., 1998; Carvelli et al., 2004).* SERT was shown to carry an uncoupled current, which is contingent on the presence of intracellular K^+^ (*Adams and DeFelice, 2003; Schicker et al., 2012*). We investigated, if such a current also exists in DAT. Accordingly, we recorded dopamine-induced currents in the presence and absence of K_in_^+^. In Fig. 4A, we show a representative current trace obtained from a DAT-expressing cell, which was evoked by the application of 30 µM dopamine in the absence of K_in_^+^. The currents recorded in the absence of K_in_^+^ were not markedly different from those evoked with physiological intracellular K^+^ concentrations (*cf*. Fig. 2A). However, when we plotted the average current amplitude and the voltage dependence thereof, we found that the currents were smaller in the absence of any K_in_^+^ (open triangles; n = 12, solid blue line in Fig. 4B) than in the presence of 160 mM K_in_^+^ (open squares; n = 12, solid black line in Fig. 4B). Differences in current amplitudes between these two conditions only reached statistical significance at potentials more positive than - 60 mV. Accordingly, we show a comparison at - 20 mV in the inset of Fig. 4B (n = 12 cells for each condition, p = 0.003).

In a recent study, we showed that SERT completed the transport cycle faster in the presence than in the absence of intracellular K^+^ (*Hasenhuetl et al., 2016*). If the same was true for DAT, it could explain the larger current amplitudes observed with physiological K_in_^+^ concentrations. We tested this notion by relying on a ‘two-pulse’ protocol (schematic representation in Fig. 4C): a saturating concentration of dopamine (i.e. 30 µM) was applied for 1 second as a first pulse. This reference pulse will force all available transporters into the transport cycle. This is followed by a second test pulse of 30 µM dopamine for 1 second. If all transporters dwell in the transport cycle, where they adopt conformations that cannot bind extracellular substrate, the test pulse will fail to evoke a peak current. However, with increasing time intervals between the reference and the test pulse, more transporters will return to the initial state and bind substrate again to produce a peak current. Accordingly, the rate of peak current recovery is a measure of the rate by which the transporter returns to the resting state (i.e. the sodium bound outward facing conformation - T_o_Na). Fig. 4C and 4D show representative original traces recorded at - 50 mV and the average time-dependent recovery of the peak current, respectively. It is evident from the time-course of peak current recovery was comparable in the presence (squares in Fig. 4D) and absence of intracellular K^+^ (circles in Fig. 4D). Thus, K_in_^+^ did not accelerate the recovery of the peak current in DAT. The tested ionic condition biases DAT into the forward direction, where the peak recovery rate becomes a measure of the rate of substrate uptake (cycle turnover rate). These data, therefore, suggest that the rate of substrate uptake is not affected by K_in_^+^. We further validated this conclusion by assessing uptake of APP^+^ in the presence and absence of intracellular K^+^: representative traces of APP^+^ uptake by DAT expressing cells, which were filled with internal solutions containing either 0 mM or 160 mM K_in_^+^ and clamped to −60 mV, are shown in the upper and lower panel, respectively, of Fig. 4E. Based on recordings from 14 cells each (Fig. 4F), it is evident that APP^+^ uptake was independent of K_in_^+^ concentrations. In Fig.4G we show the normalized slope of the APP^+^ induced rise in fluorescence in the absence of K_in_^+^ at - 90 mV, - 60 mV, - 30 mV, 0 mV and + 30 mV (open circles). For comparison we plotted the fit of the data in Fig.1E (160 mM K_in_^+^- dashed line). The two voltage dependencies are in reasonable agreement, in further support of the idea that substrate uptake is unaffected by K_in_^+^.

### K_in_^+^ binds to the inward facing conformation of DAT

Our data indicate that K_in_^+^ does not accelerate substrate uptake by DAT but gives rise to uncoupled currents, which add up to currents coupled to substrate transport. The latter finding is difficult to explain if K_in_^+^ does not interact with DAT. We tested if K_in_^+^ did indeed bind to DAT by examining the voltage-dependence of peak current amplitudes elicited by dopamine application in the presence of 160 mM K_in_^+^. Representative current traces are shown in Fig. 5A and the normalized peak current amplitudes are plotted as a function of voltage in Fig. 5B: the green solid line represents the fit of the data points to the Boltzmann equation. The slope of the current voltage relation at 160 mM K_in_^+^ was much shallower than at 0 mM K_in_^+^ (for comparison see dashed line, which corresponds to the fit extracted from Fig. 3F, i.e. with 0 mM Na_in_^+^/ 0 mM K_in_^+^ in the internal solution). The question, however, is whether the observed slope change does indeed reflect binding of K_in_^+^ to DAT. Alternatively, it may simply result from transient occupation of the inner vestibule by K_in_^+^, because upon opening of the inner vestibule the newly available space becomes occupied by water and possibly also by ions. Occupation of the inner vestibule by K_in_^+^ may suffice to produce an outwardly directed transient current and thereby diminish the voltage-dependence of the peak current. It is possible to discriminate between these two possibilities by replacing K_in_^+^ with equimolar concentration of Li_in_^+^ (i.e. 160 mM Li^+^). It is evident from the results shown in Fig. 5C that Li^+^ did not affect the slope in any appreciable way (compare orange solid line in Fig. 5C with the blue dashed line in Fig. 5B). The black and green dashed lines in Fig. 5C were taken from Fig. 3F (152 mM Na_in_^+^) and Fig. 5B (160 mM K_in_^+^), respectively, to illustrate the difference. This observation supports the argument that the slope change is indicative of specific K_in_^+^ binding to DAT and not transient occupancy of the inner vestibule of DAT by monovalent cations. In fact, K_in_^+^ binding to the inward-facing conformation of DAT provides an explanation for the observation that steady-state current amplitudes were larger in the presence than in the absence of intracellular K^+^ (see Fig. 4B). We note that the slope of the current voltage relation with high intracellular K_in_^+^ was similar to the slope observed in the presence of high intracellular Na^+^ (dashed lines in Fig.5C). This suggests that K_in_^+^ can reach the site, which Na_in_^+^ has dissociated from. We speculate that this site is the Na_2_ site.

### Expanding the two-pulse protocol to confirm the voltage-dependence of dopamine uptake by DAT

We used the “two-pulse protocol” (*cf*. Fig. 3C) under different intracellular ionic conditions and membrane voltages to obtain complementary insights into the voltage-dependence of dopamine transport through DAT. Fig. 5D shows the time course of peak current recovery determined at three potentials (i.e. - 50 mV, 0 mV, + 20 mV). These were measured in the presence of a physiological intra- and extracellular ionic gradients (160 mM K_in_^+^, 6 mM Na_in_^+^). It is evident that the rate by which DAT returned to the resting state (T_o_Na) was voltage-dependent and that it was accelerated at positive potentials. This is indicative of a lower cycle completion rate in the physiological forward transport mode and of an increased likelihood for return of the transporters to the resting state with the substrate bound via the exchange mode. We also measured the peak current recovery rate in the absence of K_in_^+^ and Na_in_^+^, at - 50 mV, 0 mV and + 20 mV: the recovery rate remained voltage-dependent, but the voltage-dependence was less pronounced than in the presence of physiological ion gradients (*cf*. Fig. 5E and 5D). The shift in voltage-dependence was not due to the absence of K_in_^+^ but rather due to the presence of 6 mM Na_in_^+^ in the experimental conditions of Fig. 5D. We confirmed this conclusion by measuring the voltage dependence with 160 mM K_in_^+^ and 0 mM Na_in_^+^ (*cf*. Fig. 5E and 5F). This observation also supports the conclusion that the faster return at positive potentials is due to transporters operating in the exchange mode; see-sawing through the exchange mode requires Na_in_^+^ (*Khoshbouei et al., 2003; Kahlig et al., 2005; Hasenhuetl et al., 2016*).

### The uptake rate of APP^+^ and dopamine is unaffected by Cl_in_^-^

In Fig.5G we assessed the effect of intracellular Cl^-^ on uptake of APP^+^ (10 µM) through DAT at - 60 mV. Shown are the slopes of the APP^+^ induced rise in fluorescence at 0.5 mM (n = 7) and 142 mM Cl_in_^-^ (n = 8), respectively. The slopes were not different (p= 0.93; Mann Whitney test) suggesting that Cl_in_^-^ plays no role in the rate of APP^+^ uptake. In a recent study, Erreger and coworkers (*Erreger et al., 2008*) measured the time course of peak recovery at 2 mM and at 140 mM Cl_in_^-^ and found the cycle rate reduced by a factor of about three at low Cl_in_^-^, when d-amphetamine was used as the substrate. If true, this implies that substrates differ in their dependence on Cl_in_^-^. In fact, when we measured the peak current recovery at 0.5 mM and 142 mM Cl_in_^-^, respectively using dopamine as the substrate we did not observe a change in recovery rate (Fig. 5H). The data points in the graph are the normalized peak current amplitudes induced by 30 µM dopamine as a function of the time interval between the reference and the test pulse. The recordings were conducted at - 50 mV. The rates estimated by fits of a mono-exponential function to the data were 2.5 s^-1^(0.5 mM Cl_in_^-^ (n = 6); open circles, blue solid line) and 3.0 s^-1^ (142 mM Cl_in_^-^ (n = 6); open squares, black solid line). Difference in these rates was not statistically significant (p=0.086, F-test). Hence, our results show that the uptake rate of APP^+^ and of dopamine are independent of Cl_in_^-^ at concentrations ranging from 0.5 mM to 142 mM. We note, however, that for technical reasons we could not test Cl_in_^-^ concentrations below 0.5 mM.

### A kinetic model for the transport cycle of DAT

DAT and SERT share many functional properties; a non-exhaustive list of common features includes: (i) their inward-facing conformations bind Na^+^ in a voltage-dependent manner (this study and *Hasenhuetl et al., 2016*); (ii) they both handle Cl^-^ similarly (*this study and Hasenhuetl et al., 2016*); (iii) the return of the transporter from the substrate-free inward-to the substrate-free outward-facing conformation is the rate-limiting step for substrate turnover by both transporters (*Erreger et al., 2008; Schicker et al., 2012*). Strikingly, however, our current observations show that all major differences between SERT and DAT can be accounted for by the difference in handling of K_in_^+^, e.g.: (i) K_in_^+^ fails to render the peak current recovery by DAT voltage-independent; (ii) K_in_^+^ has a very modest effect on the steady current amplitude and (iii) does not affect the rate of substrate transport in DAT. In all three instances, K_in_^+^ has the opposite effect on SERT (*Hasenhuetl et al., 2016*). Our current observations showed, though, that K_in_^+^ bound to DAT in a voltage-dependent manner. Thus, electrogenic binding of K^+^ is another common feature in SERT and DAT. It is, however, enigmatic why K^+^_in_ binding to both transporters affects the function of these transporters so differently. We hypothesized that the differences between SERT and DAT can all be explained by positing that, in the return step from the substrate-free inward-to the substrate-free outward-facing conformation, K_in_^+^ is counter-transported by SERT, but left inside by DAT. We tested the plausibility of this conjecture by resorting to kinetic modelling. As a starting point, we used the previously proposed kinetic model of DAT by Erreger and coworkers (*Erreger et al., 2008*). The salient features of this model were: (i) only one Na^+^ ion is co-transported with the substrate, (ii) the return from the substrate free inward-to the substrate-free outward-facing conformation (the return step) is not contingent on prior Cl^-^ dissociation and (iii) the return step is rate limiting for substrate transport. The refined model in Fig. 6A conforms to all of the above. This is evidenced by the fact that the original scheme (shaded in grey) is nested within our proposed model. The notion that only one Na^+^ is co-transported with the substrate is widely accepted for SERT (*Rudnick, 1977*). For DAT, it has been proposed that two Na^+^ ions are co-transported (*Krueger, 1990; McElvain and Schenk, 1992*). However, we adhered to the stoichiometry posited by Erreger *et al.* because our experimental results support the idea that only one Na^+^ ion is co-transported in each cycle (see Fig. 3F). Our data show that Na^+^ binding to the inward facing conformation of DAT is a voltage-dependent process. We incorporated this into the model by assigning a valence of 1 to this reaction.

**Fig. 6.**
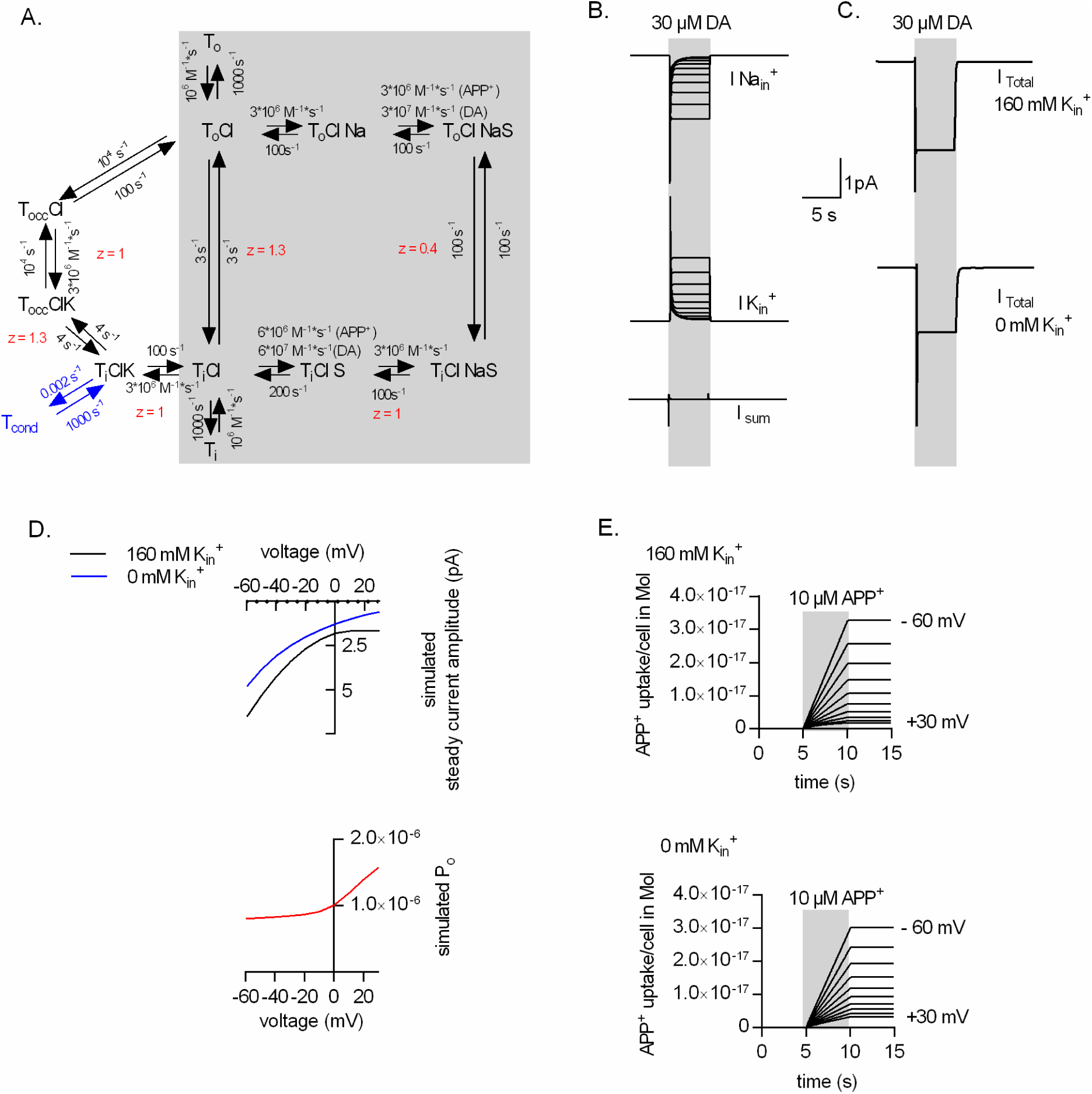
A kinetic model of the transport cycle of DAT. **A.** Schematic representation of the transport cycle. Shaded in grey is the original scheme proposed by Erreger et al. (*2008*), which is nested in our refined scheme. The letter T stands for transporter, the subscripts o, i, occ and cond stand for outward facing inward facing, occluded and conducting state, respectively. The annex Cl, Na, K indicate when the corresponding ion is bound; the annex S when a substrate is bound. We parameterized substrate binding by using different sets of rate constants for APP^+^ and dopamine. The z values indicated in red are the valences, which are ascribed to 6 partial reactions. All other reactions were assumed to be voltage-independent. Indicated in blue is the conducting state, which was modeled as a selective pore for Na^+^ ions. **B.** The upper panel, shows the simulated current component produced by Na^+^ ions dissociating into the cytosol (calculated for voltages ranging from - 60 mV to + 30 mV). The panel in the middle shows the current component produced by incoming K^+^ ions (also for voltages ranging from - 60 mV to + 30 mV). The lower panel shows the sum of the two. It is evident that the two currents cancel each other out. **C.** Simulated currents induced by dopamine (30 µM) in the presence and absence of 160 mM K_in_^+^. **D.** Voltage-dependence of the steady current component with and without 160 mM K_in_^+^ (upper panel). In the lower panel we show the open-probability of the conducting state as a function of voltage. The conducting state is gated by voltage. **E.** Simulated uptake of 10 µM APP^+^ in the presence and absence of 160 mM K_in_^+^ in the upper and lower panel, respectively. Uptake was simulated for voltages ranging from - 60 mV to + 30 mV. There is little effect of K_in_^+^ on APP^+^ uptake.

The refined model takes into account the action of K_in_^+^ on DAT, which was not explored earlier (*Erreger et al., 2008*). We assigned a valence of 1 to the reaction of K_in_^+^ binding to the inward facing conformation of DAT. This assured that the inwardly directed current, produced by outgoing Na_in_^+^ ions dissociating into the cytosol, was canceled out by the outwardly directed current generated by the incoming K_in_^+^ ions. This point is illustrated in Fig. 6B, where we show the simulated current component produced by dissociating Na_in_^+^ ions (upper panel) and associating K_in_^+^ ions (middle panel) individually for voltages ranging from - 60 mV to + 30 mV. When summed, these currents canceled each other out (see lower panel). In the refined model, we accounted for the uncoupled current component observed in the presence of K_in_^+^ by adding a conducting state, which is in equilibrium with the K_in_^+^-bound inward-facing conformation (indicated in blue). This is akin to how we previously modeled the uncoupled current through SERT. In contrast to SERT, where compelling evidence exists that K^+^ is antiported, we assumed that K^+^ is shed off by DAT in the return step from T_i_ to T_o_, i.e. the substrate-free inward-to the substrate-free outward-facing conformation. It was necessary to subdivide this transition into three consecutive reactions to incorporate this concept into the kinetic model: in the first reaction (when viewed in the clockwise direction), DAT adopts an inward-facing conformation on the trajectory to the occluded state, to which K_in_^+^ can still bind, but with reduced affinity. In the second reaction, DAT fully occludes after shedding off K_in_^+^. Following occlusion, the transporter can then rearrange and adopt the outward-facing conformation. In Fig. 6C, we show synthetic traces of dopamine-induced currents in the assumed presence (upper panel) and absence (lower panel) of K_in_^+^ at - 60 mV, as predicted by the model. The amplitude of the steady current in the simulation was larger with K_in_^+^ present. The upper panel of Fig. 6D displays the calculated voltage-dependence of the steady current with and without K^+^_in_: the synthetic data are in reasonable agreement with the data shown in Fig. 4B. The lower panel of Fig. 6D shows the simulated open probability (P_o_) of the conducting state as a function of voltage. It is obvious from this plot that the conducting state is gated by voltage. This property can be readily explained: T_icond_ is assumed to be in equilibrium with T_i_ClK. Occupancy of T_i_ClK is increased at positive potentials, due to the voltage dependent slowing of the substrate-free return step. This may explain the parallel shift in the two current-voltage relations, i.e. P_o_ is lower at negative potential but the driving force for Na^+^ is larger; the opposite situation is seen at positive potentials.

Our kinetic model of DAT was also able to faithfully predict cellular uptake of APP^+^. In the model, we accounted for the larger K_M_ value observed with APP^+^ by assuming a 10-fold lower association rate for APP^+^ than for dopamine. It is worth pointing out that a lower association rate can also explain why we failed to see an APP^+^-induced peak current at concentrations below 30 µM (Fig. 2B). In Fig. 6E, we simulated uptake of APP^+^ in the presence and absence of K_in_^+^ for voltages between - 60 mV and + 30 mV: it is evident that uptake of APP^+^ and the voltage dependence thereof was only affected to very less extent by the absence of intracellular K^+^.

Our modeling exercise demonstrates that DAT can bind K_in_^+^ but does not antiport it, i.e. it is possible to implement this concept into a kinetic model without violating microscopic reversibility. More importantly, we show that the assumed inability of DAT to countertransport K_in_^+^ can explain (i) the presence of a coupled current component and (ii) the lack of effect of K^+^_in_ on the voltage-dependence of substrate transport. We stress that these two observations would have been difficult to explain, if K^+^ was antiported by DAT.

## Discussion

Mechanistic and functional insights into the mode of operation of monoamine transporters have been derived either by relying on radioligands, which bind to monoamine transporters and are substrates or inhibitors, or by electrophysiological recordings of transport-associated ion fluxes. Radioligand-based techniques, used to assess global transporter-mediated uptake or release, lack membrane voltage control, temporal resolution and sampling frequencies to resolve individual partial reactions that constitute a transport cycle. These issues can be addressed by single cell electrophysiological recordings. These measurements, however, rely on transporter-mediated currents as an indirect surrogate for substrate uptake and are often confounded by additional conductance not directly related to substrate transport (i.e. uncoupled currents). Fluorescent substrates of monoamine transporters, on the other hand, help circumvent limitations imposed by radioligand assays or electrophysiological recordings by monitoring transporter-mediated uptake in real time (*Schwartz et al., 2003; Mason et al., 2005; Schwartz et al., 2005; Solis Jr. et al., 2012; Karpowicz Jr et al., 2013; Wilson et al., 2014; Zwartsen et al., 2017*). In this study, we investigated the transport cycle of DAT, in single DAT-expressing HEK293 cells, by employing simultaneous measurements of substrate-induced currents recorded in the whole-cell patch-clamp configuration and cellular uptake of the fluorescent substrate APP^+^. The key finding of our study is that the voltage-dependence of DAT-mediated substrate uptake and ionic currents is dictated by intracellular binding of K_in_^+^ to DAT. This conclusion is based on the following observations: (i) both, uptake of APP^+^ and substrate-induced currents decreased at positive voltages in DAT-expressing cells. (ii) Replacing K _in_^+^ with the inert cation NMDG^+^ did not influence the rate of substrate uptake but led to reduction in amplitudes of substrate-induced DAT steady-state currents. (iii) The current-voltage relationship of peak currents was identical when patched with internal solutions, which contained either high K_in_^+^ or Na_in_^+^ concentrations, indicating mutually exclusive binding of the two ions on the same site (presumably Na_2_). This relationship was steeper when Na_in_^+^ or K_in_^+^ were replaced intracellularly by either NMDG^+^ or Li^+^. (iv) A kinetic model can simulate K_in_^+^ binding to DAT as transient and reversible, without requiring antiport of K_in_^+^ when the transporter isomerizes from T_i_ to T_o_, the empty inward open to the outward open apo-state. This model accounts for all experimental observations, which support the conclusion that the transient K_in_^+^ binding to DAT is electrogenic, tempers the electrogenic dissociation of Na_in_^+^ from DAT to a certain extent (but not completely) and gives rise to uncoupled currents (that is, currents unrelated to the DAT transport cycle).

Conclusions made from earlier studies assessing voltage dependence of DAT mediated uptake were conflicting (*Hoffman et al., 1999; Prasad and Amara, 2001*). We addressed these discrepancies by studying DAT mediated uptake of APP^+^. APP^+^ is a fluorescent analog of the neurotoxin MPP^+^, a known substrate for all 3 monoamine transporters (*Scholze et al., 2002*). APP^+^ fluoresces in non-polar environments but is quenched in aqueous solutions (*Karpowicz Jr et al., 2013*). Upon transporter-mediated entry into cells, APP^+^ binds to membranes of organelles (mostly mitochondria) due to the permanent positive charge on APP^+^. This leads to APP^+^ sequestration in cells expressing monoamine transporters and low levels of cytosolic APP^+^. Therefore, cellular APP^+^ uptake (but not release) can be directly monitored in real time by recording the rise of fluorescence resulting from intracellular sequestration. We obtained recordings of the time course by which APP^+^ accumulated in cells expressing DAT. The cells were measured in the whole-cell patch-clamp configuration, which allowed us to directly assess the effect of voltage on this process. We were able to demonstrate that APP^+^ uptake by DAT is voltage-dependent, with reduced uptake at increasingly positive voltages. To ensure that the conclusions obtained from our recordings of APP^+^ uptake also hold true for uptake of dopamine (which does not fluoresce), we analyzed currents induced by APP^+^. As shown by us and others (*Solis Jr. et al., 2012*), APP^+^-induced currents showed the hallmark features of those induced by other DAT substrates such as dopamine or D-amphetamine (*Erreger et al., 2008*), an inwardly directed peak current and a steady current component. Finally, the K_M_ values for APP^+^ were identical, when determined by the substrate-induced currents and by examining the concentration-dependent slope of the APP^+^-induced rise in fluorescence.

Another aspect to our study was to understand why the structurally similar SERT and DAT differ so much in transport kinetics and handling of co-substrate binding manifested as ionic stoichiometry. In a previous study, we explored the transport cycle of SERT in great detail to generate a comprehensive model of the transport cycle of SERT (*Hasenhuetl et al., 2016*). It is evident that DAT resembles SERT in most aspects. We show here that all major differences can be accounted for by the distinct handling of K_in_^+^: in SERT, physiological K_in_^+^ concentrations accelerate the rate of substrate uptake (i.e. peak current recovery rate) such that it is 2-fold faster than in the absence K_in_^+^. In DAT, K_in_^+^ did not influence rate of substrate uptake. In addition, in the presence of physiological K_in_^+^ concentrations, the cycle completion rate of SERT is independent of voltage. This was not the case in DAT. In both transporters, release of Na^+^ from the inward facing conformation is electrogenic. In SERT, this electrogenic Na_in_^+^ dissociation is canceled out by electrogenic K_in_^+^ binding to the inward-open empty transporter, thereby rendering the cycle completion rate voltage-independent. In DAT, however, the cycle completion rate remained voltage-dependent despite the fact that K^+^_in_ also bound in a voltage-dependent manner. K_in_^+^ is also relevant to account for the distinct nature of the steady-state current component in SERT and DAT. The steady current component carried by the electroneutral SERT is produced by an uncoupled Na^+^ flux through a channel state that is in equilibrium with the K_in_^+^-bound inward-facing conformation (*Schicker et al., 2012*). Replacing K_in_^+^ with NMDG^+^ completely abrogates SERT mediated steady-state currents. DAT-mediated transport is accompanied by the translocation of coupled net positive charges in each cycle. Thus, DAT-mediated steady-state currents were originally modeled to be strictly coupled to substrate transport (*Erreger et al., 2008*). However, our data suggests that DAT also carries an uncoupled current component, which adds up to coupled currents associated with DAT-mediated ionic transport. This uncoupled current in DAT, just like in SERT, is contingent on the presence of intracellular K_in_^+^. Binding of K_in_^+^ to DAT at the Na_2_ site was proposed earlier by a study that employed extensive molecular dynamic simulations to understand Na^+^ dissociation intracellularly by DAT (*Razavi et al., 2017*). Our results, which showcase similarities in voltage dependencies of peak amplitudes in the presence of either high Na_in_^+^ or high K_in_^+^ (Fig. 5C), are in agreement of this hypothesis.

The most parsimonious explanation for all differences between SERT and DAT was to posit that K^+^ is antiported by SERT but not by DAT. Accordingly, our analyses provide a unifying concept of substrate transport through SERT and DAT, i.e., SERT and DAT are equivalent in all aspects of their transport cycle but one: in SERT the binding site for K_in_^+^ remains intact upon conversion of the transporter from the inward to the outward facing conformation. In contrast, this binding site is less stable in DAT. The resulting loss in affinity leads to the shedding of K_in_^+^ prior to the return step. The repercussions of this subtle difference are profound: SERT and DAT differ (i) in their voltage-dependence of substrate uptake, (ii) in the nature of the substrate-induced current and (iii) in the energy sources exploited for concentrative substrate transport. If DAT does not antiport K_in_^+^, its concentrative power must be independent of the existing K^+^ gradient. On the other hand, if the stoichiometry of DAT is electrogenic, a change in membrane voltage is predicted to increase or decrease substrate uptake at steady state depending on the direction of the voltage change. In this context, it is important to note that the experiments conducted in the present study all report on substrate transport at pre-steady state. Additional insights on whether or not DAT can antiport K_in_^+^ can come from experiments conducted at the thermodynamic equilibrium. Such experiments need to be performed by, for instance, employing a vesicular membrane preparation that contains reconstituted DAT. Such preparations allow for the control of the inner and outer ion composition and membrane voltage while preventing the substrate to escape from the vesicular confinement. However steady-state assessment of transporter mediated substrate uptake is hindered by the fact that DAT and SERT can also transport substrate in the absence of K_in_^+^. These observations are difficult to reconcile with the concept of transport by fixed stoichiometry. We, therefore, surmise that SERT and DAT, operate with a mixed stoichiometry. Based on our data we conclude that DAT is less likely than SERT to antiport K^+^, because we cannot rule out that DAT can occasionally carry the K^+^ ion through the membrane. Conversely SERT antiports K^+^ in the majority of its cycles but may return empty in some instances. We thus believe that the differences between these two transporters in regard to their handling of K_in_^+^ represents a continuum, as opposed to divergence, in ionic coupling and kinetic decision points during substrate transport. The difference between SERT and DAT represent different solutions to an inherent trade-off and may reflect an adaptation to physiological requirements: because of electrogenic binding and subsequent counter-transport of K^+^, SERT operates in the forward transport mode with a constant rate regardless of membrane potential, but it cannot exploit the membrane potential to fuel its concentrative power. In contrast, DAT can harvest the energy of the transmembrane potential to fuel its concentrative power. As a trade-off, the substrate uptake rate DAT is voltage-dependent and strongly reduced upon membrane depolarization.

## Acknowledgments

We thank Verena Burtscher for discussion and comments on the data. This work was supported by the Austrian Science Fund (FWF) grant P 31599 and P 31813 to W.S, W1232 to H.H.S and the Vienna Science and Technology Fund (wwtf) grant CS15-033 to H.H.S and LS17-026 to M.F.

